# Shear Stress Modulates Endothelial Ca^2+^ Signaling and Barrier Integrity in a Microfluidic Organ-on-a-Chip Platform

**DOI:** 10.1101/2025.10.30.685554

**Authors:** Ludovica Montesi, Mattia Tiboni, Mario Hagel, Luca Casettari, Rossana Rauti

## Abstract

Endothelial cells (ECs) line the blood vessels and form the primary barrier between the bloodstream and the brain. Blood flow exerts a modulatory effect on the endothelial phenotype, and evidence indicates that capillary-like fluid shear stress enhances endothelial tight junctions and transporters. ECs possess mechanosensitive channels that activate endothelial responses and modulate their functions. Among the various responses to shear stress, morphological adaptations have been the most extensively studied, while functional live responses remain mostly unexplored. Calcium has been identified as key modulators that translates mechanical stimuli into biological processes, regulating endothelial activity. In this study we investigate the effect of acute and long-term shear stress on endothelial cells through live calcium imaging and immunocytochemistry, by using a modular 3D printed organ-on-a-chip, capable of simulating *in-vivo* capillary and enabling the possibility to study cellular crosstalk.

## 1. Introduction

Human endothelial cells line the blood vessels intima, and represent the main interface between the circulating blood and the brain parenchyma (Maurya et al., 2021). Endothelial cells play a key role in maintaining the integrity of the blood–brain barrier (BBB), regulating molecular exchange, and preserving neural homeostasis. Accumulating evidence suggests that Fluid Shear Stress (FSS), the tangential force applied to the vascular endothelium generated by the blood flow, play an important role in modulating vascular endothelium phenotype (Wang et al., 2021).

Under physiological, laminar flow conditions, FSS supports endothelial stability by promoting junctions’ organization, maintaining a quiescent phenotype, and suppressing inflammatory activation. In contrast, disturbed or oscillatory flow patterns disrupt junctional integrity and lead to both morphological and functional barrier alterations (García-Polite et al., 2017; Maurya et al., 2021; Nunez et al., 2024).

The ability of endothelial cells to sense and convert these mechanical cues into biochemical signals is known as mechanotransduction. This process involves the activation of membrane and cytoskeletal complexes that initiate intracellular cascades, ultimately modulating gene expression and barrier behavior. Several studies have shown that shear stress can influence the organization of cell–cell junctions such as VE-cadherin, thereby affecting endothelial permeability and overall vascular function (Jackson et al., 2023; Bathrinarayanan et al., 2025).

Most existing research on shear stress has focused on morphological changes or the expression of structural and molecular markers (Cronin et al., 2025; Franz et al., 2025). However, functional aspects, such as real-time intracellular signaling, are still not well understood. In particular, calcium dynamics play a pivotal role in endothelial mechanotransduction, as transient Ca^2+^ elevations regulate a wide range of processes including cytoskeletal remodeling, nitric oxide synthesis, and permeability control (Charbonier, et al., 2019; Hong et al., 2024; Peng et al.,2023). Despite its importance, the precise mechanisms driving Ca^2+^ mobilization in response to different flow regimes remain unclear. Both Ca^2+^ entry through mechanosensitive channels and Ca^2+^ release from intracellular stores, such as the endoplasmic reticulum, have been implicated, but their relative contributions under distinct shear stress conditions are still debated (Nunez et al., 2024).

A major obstacle to investigating these processes has been the lack of experimental systems capable of combining *in-vitro* endothelial culture with controlled flow exposure. The emergence of Organ-on-a-Chip (OoC) technologies has opened new perspectives in this field, providing engineered platforms that integrate human-derived tissues within perfusable microchannels to mimic physiological environments (Ingber, 2022). Nevertheless, many existing devices are limited by low imaging compatibility, which restricts their application for live-cell calcium imaging and functional assays (Rauti et al., 2021).

In the present work, we introduce a 3D-printed, modular Organ-on-a-Chip platform specifically designed to study the effects of shear stress on human endothelial cells. The system enables precise control of flow conditions, allowing the simulation of both physiological and pathological shear stress levels. Its architecture supports long-term cell culture, real-time calcium imaging, and subsequent immunofluorescence analysis, offering a versatile approach to investigate how mechanical forces influence endothelial signaling, barrier integrity, and overall BBB function.

## 2. Materials and Methods

### 2.1 Chip Design and Fabrication

The chip design was adapted from a previous publication by our research group (Rauti et al., 2021; Rauti et al, 2023), where a dedicated channel was incorporated to achieve improved control over shear stress and flow conditions. The chip was designed using CAD Autodesk Fusion 360 and exported as single file. A schematic representation of the chip can be visualized in **Figure 1**. Prior to printing, model surfaces were checked, and supports were added using PreForm software (PreForm 3.0.1, Formlabs, Inc.). Then, the chips were printed in a stereolithography Form 3B+ printer (Formlabs, Somerville, Massachusetts), using a biocompatible, transparent resin (BioMedClear V1, Formlabs), with unique mechanical and optical properties. After printing, the chips were washed in isopropyl alcohol (Avantor), to remove the unreacted resin, and then cured and dried in a UV curing system (Formlabs).

**Figure 1.**
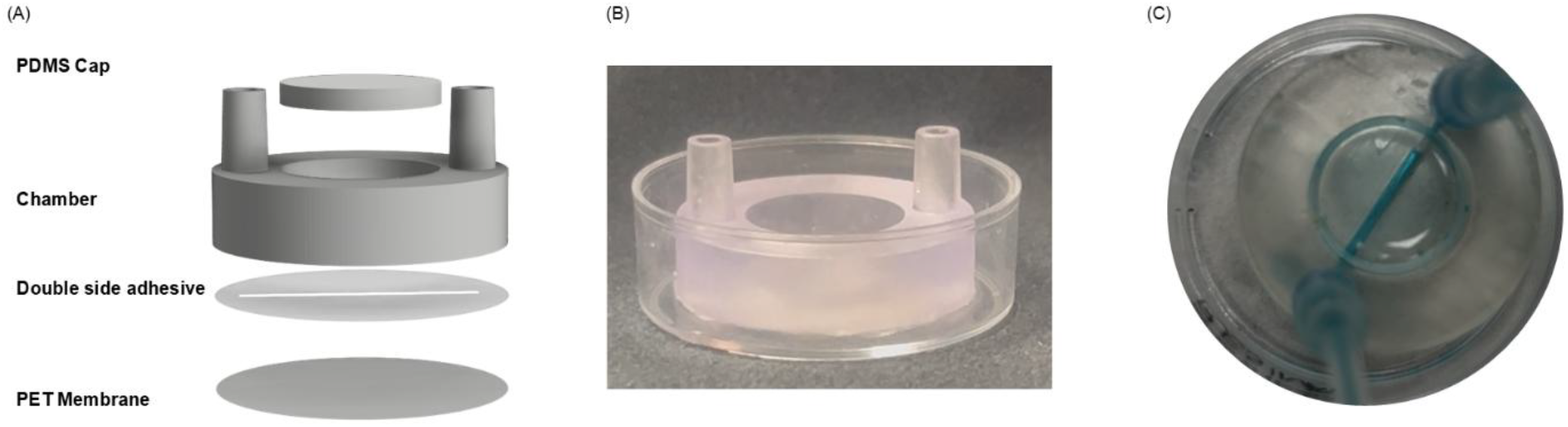
Open-top chip design. (A) Schematic exploded view of the chip composed of i) PET porous membrane, ii) double adhesive tape, iii) resin module with channel, iv) PDMS cap. (B) Photograph of the chip. (C) P Photograph of the chip with channel perfused with blue food dye.

#### Fabrication of additional components

We used CAD Autodesk Fusion 360 software to design the master-mold for fabrication of the device’s additional component, the Polydimethylsiloxane (PDMS) cap. The molds were printed with PreForm software (PreForm 3.0.1, Formlabs, Inc.). Then, the molds were filled with PDMS prepared by mixing Sylgard 184® (Dow Corning, Midland, MI) with the curing agent at a ratio of 1:10, followed by curing at 60 °C overnight. The resulted PDMS caps were cleaned in ethanol, dried at room temperature and sterilized 30 min with UV lamp.

Polyethylene (PET) membranes (0.4 μm pore size, it4ip S.A., Belgium), 12 μm thick, were cut to size with their protective backing on. The protective backings were then removed, and the PET membranes were attached to the chip through a double side adhesive (3M™ 467MP) on the bottom (a detailed description of the chip assembly will be provided in the Results section).

#### Chip Sterilization

The ready-to-use assembled chip was sterilized using ethanol for 1h, under the laminar flow hood, rinsed with MilliQ water for three times, and then exposed to UV lamp for 1 hour.

For the flow condition, the tubing was sterilized by perfusing ethanol throughout the system at a flow rate of 200 µL/min for 30 min, to ensure a proper sterilization of the system. Following that, PBS was flushed into the entire system for an additional 30 min at the same flow rate, to ensure the complete removal of ethanol.

#### Flow gradients inside the chip

Flow was controlled by an external peristaltic pump (ISM931 V4.00, Ismatec, Cole-Parmer GmbH, Werthein, Germany), and connections were in Tygon® tubing. The input tube was connected to the inlet of the chip, and the output was connected to a reservoir via the peristaltic pump.

### 2.2 Computational fluid dynamics (CFD)

Computational fluid dynamics (CFD) simulations were performed using ANSYS Multiphysics to investigate the wall shear stress and velocity field within a custom-designed microfluidic chip. The ANSYS Fluid Flow solver was employed to simulate flow conditions relevant for cell culture at the chip’s bottom membrane, where cells are seeded.

#### Chip Geometry and Mesh Generation

The chip geometry was modeled according to experimental design specifications, featuring a cylindrical chamber with inlet and outlet ports of 2 mm diameter (cross-sectional area: 3.14×10-6m^2^). The interface between the chip walls and fluidic domain was defined as a static boundary, while the bottom membrane was treated as a rigid, non-deformable surface representing the cell culture substrate. For mesh generation, a hybrid approach was utilized—hexahedral elements for rectangular portions and tetrahedral elements for cylindrical regions—resulting in 263,468 nodes and 786,944 elements. Inflation layers and mesh transitions were software-controlled for optimal resolution of boundary gradients.

#### Simulation Setup and Assumptions

Simulations were conducted under steady-state conditions with the following assumptions and settings:

- Incompressible, Newtonian fluid, with constant viscosity and density properties analogous to cell culture medium (water at room temperature).
- Laminar flow was prescribed due to low Reynolds numbers typical for microscale channels (<2000).
- No-slip boundary conditions at all channel walls.
- Rigid channel walls and negligible gravity or temperature effects.
- Outlet boundary was held at zero-gauge pressure (atmospheric), with prevention of reverse flow.
- Fluid domain was modeled as a closed system.

#### Data Analysis and Visualization

Wall shear stress and velocity distributions were extracted from the simulation data for the YZ cross-section and XY basal plane. Shear stress maps were visualized in dyne/cm^2^ to facilitate interpretation for biological applications. Flow profiles and stress gradients were quantitatively analyzed to relate simulated mechanical cues to physiological conditions experienced by seeded cells. All visualizations and results were generated using ANSYS output tools and secondary analysis in standard data plotting software.

### 2.3 Cell culture

#### Endothelial culture

For the endothelial model, Human Umbilical Vein Endothelial Cells (HUVEC, PromoCell) were used. After thawing, the HUVECs were expanded in low-serum endothelial cell growth medium (PromoCell), at 37 °C with 5% CO_2_ in a humidifying incubator and used at passage p2-p4. Cells were grown at 80-90% confluence before being seeded inside the device. Before seeding, the PET membrane was treated with Entactin-Collagen IV-Laminin (ECL) Cell Attachment Matrix (Merck) diluted in Dulbecco’s Modified Eagle Medium (DMEM; 10 μg/cm^2^) and incubated for 1h at 37 °C with 5% CO_2_. Then, the HUVECs, harvested using trypsin/EDTA solution, were seeded inside the chip at a density of 20.000/cm^2^ and grown for 4-5 days.

For the 24h flow exposure, the tubing was cleaned and sterilized as described above. After seeding the cells, the entire system (chip, tubing, and peristaltic pump) was placed in the incubator, and the peristaltic pump was activated to perfuse culture medium at a constant flow rate of 118µL/min.

### 2.4 Live Calcium Imaging

For Calcium (Ca^2+^) imaging experiments, cell cultures were loaded with the cell-permeable Ca^2+^ dye Oregon Green 488 BAPTA-1 AM (Life Technologies). Cultures were incubated for 45 minutes at 37° with 5% CO_2._ Subsequently the media was rinsed, and replaced with fresh endothelial media. The chip containing the Oregon Green loaded cultures was placed on an inverted microscope (Nikon Eclipse Ti2) and observed with a 20x objective (0.50 NA) using a DS-10 Camera (Nikon). Images were acquired every 250 ms, for 5 minutes, under continuous illumination. The Ca^2+^ dye was excited at 488 nm using appropriate filters cube set and a LED lamp (Nikon). Video analysis was done using the open-source imaging software Fiji (Imagej; Schindelin et al., 2012), and Clampfit software (pClamp suite, 10.2 version; Axon Instruments)

### 2.5 Immunocytochemistry

HUVECs cells were rinsed in Phosphate Buffered Saline (PBS) and fixed in 4% paraformaldehyde (PFA) for 20 min at Room Temperature (RT). Immunocytochemistry was carried out after permeabilization with 0.1% Triton X-100 (Sigma-Aldrich) in PBS for 10 minutes RT and blocking for 30 min with 5 % FBS in PBS. The primary antibody, anti VE-cadherin (1:200; Cell Signaling) was applied overnight in PBS and 5% FBS, at 4 °C. Cells were then washed in PBS three times and then incubated with the secondary antibody, anti-rabbit Alexa Fluor-594 (1:500; Invitrogen) in 5% FBS in PBS for 2 hours at RT. After two washes with PBS, cells were incubated with DAPI (1:10.000 in PBS; Cell Signaling) for 10 min at RT to stain the nuclei. After two washes in PBS, imaging was carried out using an inverted epifluorescence microscope (Nikon Eclipse Ti2), with appropriate filter cubes and equipped with 20x/0.50 NA and 40x/0.75 NA objectives. Image reconstruction and processing were done using the open-source imaging software Fiji (Imagej; Schindelin et al., 2012).

### 2.6 Statistical analysis

The results are presented as the mean ± standard error (SEM). All data were statistically analyzed using GraphPad Prism 7.0 software (GraphPad Software, SD, USA). The comparison of means between groups was performed by Fisher’s exact test or t-test. A statistically significant difference between two data sets was assessed and P < 0.05 was considered statistically significant.

## 3. Results and Discussion

In this work, we explored the effect of shear stress on endothelial cells through an open-top 3D printed OoC, capable of sustaining long-term culture, and allowing the modulation of different flow rates in a controlled environment.

### 3.1 Endothelial Layer Formation of the 3D-Printed Chip

Most Organs-on-a-Chip or microfluidic devices are fabricated from PDMS, which is biocompatible, transparent, and has good gas permeability. However, a major limitation of PDMS is its hydrophobicity, which causes substantial absorption of hydrophilic materials, potentially altering the microenvironment and experimental outcomes (Rauti et al., 2021). Moreover, in some cases, chip fabrication requires specific know-how and facilities (Rauti et al., 2021; Chen et al., 2022; Mehta et al., 2022). To overcome these challenges, our microfluidic chip was made by stereolithography 3D-printing, using a biomedical clear resin (see Materials and Methods), chosen for its optical transparency and biocompatibility. The use of 3D-printing enables the design of the desired platform to be quickly modified, and it reduces the need for multi-step fabrication needed in “standard” Organs-on-a-Chip. Furthermore, the use of 3D-printing reduces the fabrication time of the Organ-on-a-Chip from several days to a few hours, as well as the possibility to use not-absorbing materials (Rauti et al., 2021). Furthermore, our design also facilitates cell seeding, medium exchange, and direct microscopic observation, making it particularly suitable for long-term live-cell imaging under controlled shear stress conditions.

The chip design was based on a prior publication by our research group (Rauti et al., 2021; Rauti et al, 2023), adapted with a dedicated channel to specifically control shear stress and flow conditions. The chip is composed of a main cell culture chamber with an external diameter of 28 mm and an inner diameter of 15 mm, which hosts a channel, with dimensions 2 mm height, 1.2 mm width, and 18 mm length. Inlet and outlet channels on the upper part of the chip enable the chamber to be connected to a peristaltic pump, which allows a precise control of the fluid flow **(Figure 1**); the inlet and outlet channels are 1 mm high and 1mm long, with external diameter of 4 mm. On the bottom of the resin module, a permeable and transparent polyethylene (PET) membrane (0.4 μm pore size), was sealed through biocompatible (Winkler et al., 2022) double adhesive tape **(Figure 1A, 1B)**, facilitating cell visualization and monitoring under optical and epifluorescence microscope.

To shed light on the vascular response to shear stress, we used the microfluidic chip we developed. This configuration allows having a laminar flow directly on the cells surface, close resembling the laminar shear stress on *in-vivo* condition **(Figure 1C)**. Furthermore, the close position of the cells to the bottom of the chip allowed us to perform real-time analysis, examining essential parameters such as calcium dynamics and changes in adherens junctions, which are essential functions of the endothelium serving as a barrier.

### 3.2 Computational simulation

Computational fluid dynamics (CFD) simulations were performed to characterize the fluid behavior and wall shear stress distribution within the microfluidic chip geometry using ANSYS Multiphysics and the Fluid Flow solver (Ansys Student, ANSYS Inc.). Simulations focused on the planar surface at the bottom of the system, which represents the cell-seeding site, as well as cross-sectional and volumetric flow characteristics at various inlet flow rates.

#### Shear Stress Profiles

Shear stress distribution was evaluated for volumetric flow rates of 31, 118, 150, and 500 µL/min **(Figure 2)**, calculated based on flow rates that were not damaging the cells. At all simulated flow rates, the wall shear stress was highest at the bottom plane directly adjacent to the main flow path, with pronounced uniformity in the central regions and sharp decline toward the edges, consistent with boundary effects and the no-slip condition at the walls.

**Figure 2.**
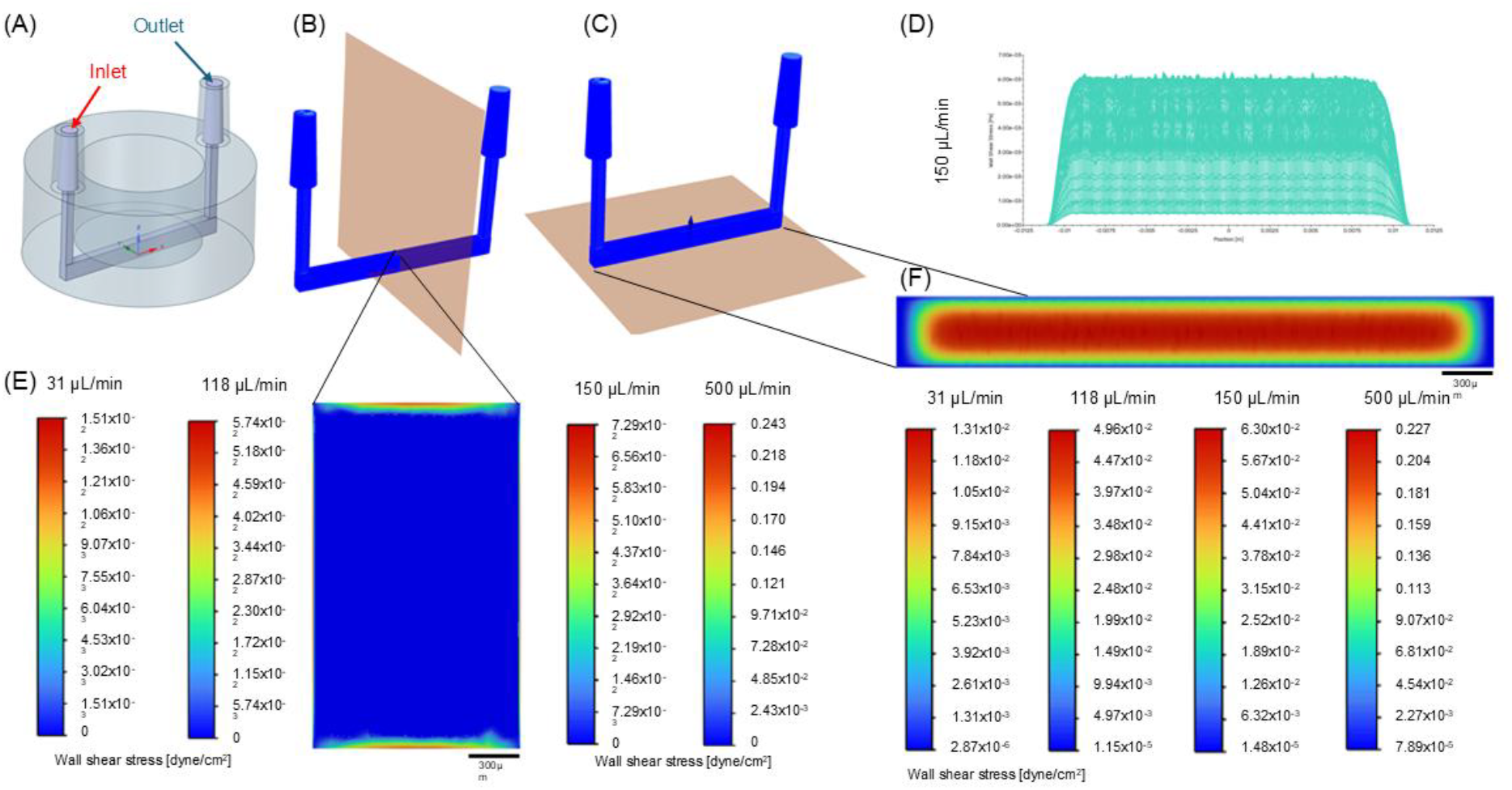
Computational fluid dynamics simulation of shear stress distribution within the microfluidic chip. (A) 3D schematic of the chip showing inlet (red) and outlet (blue) ports. (B) YZ-plane cross-section highlighting the location of the fluidic volume. (C) XY-plane view at the bottom surface of the channel where cells are seeded. (D) Wall shear stress profile along the seeded plane at 150 µL/min, illustrating the range of shear stress experienced by cells. (E) Color maps of wall shear stress (dyne/cm^2^) in the YZ-plane across different volumetric flow rates (31, 118, 150, and 500 µL/min), with corresponding quantitative scales. (F) Wall shear stress distribution along the bottom plane for each flow rate, demonstrating increased maximal shear stress with rising flow. Scale bars: 300 µm. (Images used courtesy of ANSYS, Inc.)

At the lowest flow rate (31 µL/min), the maximum wall shear stress reached approximately 1.51×10^-2^dyne/cm^2^, while increased flow rates (118 µL/min and 150 µL/min) resulted in elevated maxima of up to 5.74×10^-2^dyne/cm^2^. At 500 µL/min, the maximum shear stress further increased to 0.243 dyne/cm^2^. Across all conditions, shear stress remained relatively homogeneous over the central region of the seeded bottom plane, with small edge effects visible at lateral boundaries.

The simulations revealed that the highest wall shear stress concentrations were consistently localized to the central region beneath the primary flow pathway, with values increasing proportionally to the applied flow rate. At lower flow rates, the stress profile remains relatively uniform with a gradual drop-off toward the channel edges, whereas higher flow rates amplify both the maximum magnitude and the extent of the high-shear zone. The consistency and reproducibility of these profiles across all tested conditions underscore the robust laminar flow regime within the chip and confirm the suitability of this system for controlled under-flow measurements. These results directly inform the expected physical environment experienced by cells seeded at the channel base, enabling precise tuning of biomechanical cues through flow modulation.

#### Influence of Mesh and Boundary Conditions

The hybrid mesh employed (hexahedral in rectangular sections, tetrahedral elsewhere) and the use of a smooth inflation transition enabled accurate resolution of shear gradients at the wall. No-slip, steady-state, and incompressibility assumptions resulted in anticipated parabolic flow and stress profiles, with negligible perturbations due to gravity or temperature.

Endothelial cells were seeded in the channel, on the membrane, and interfaced to different flow rates.

### 3.3 Endothelial cells respond to shear stress with flow-dependent alterations in Ca^2+^ dynamics

To evaluate the effect of shear stress on endothelial cells we employed calcium imaging measurements (**Figure 3**). This approach allowed us to evaluate the calcium transients in cells population at single-cell resolution. After culturing endothelial cells for 4-5 days in the microfluidic chip, they were stained with Oregon Green 488 BAPTA-1 AM (membrane-permeable Ca^2+^ dye). To precisely control flow rate and shear stress, the chip was connected to a peristaltic pump and placed under the epifluorescence microscope **(Figure 3A and 3B)**.

**Figure 3.**
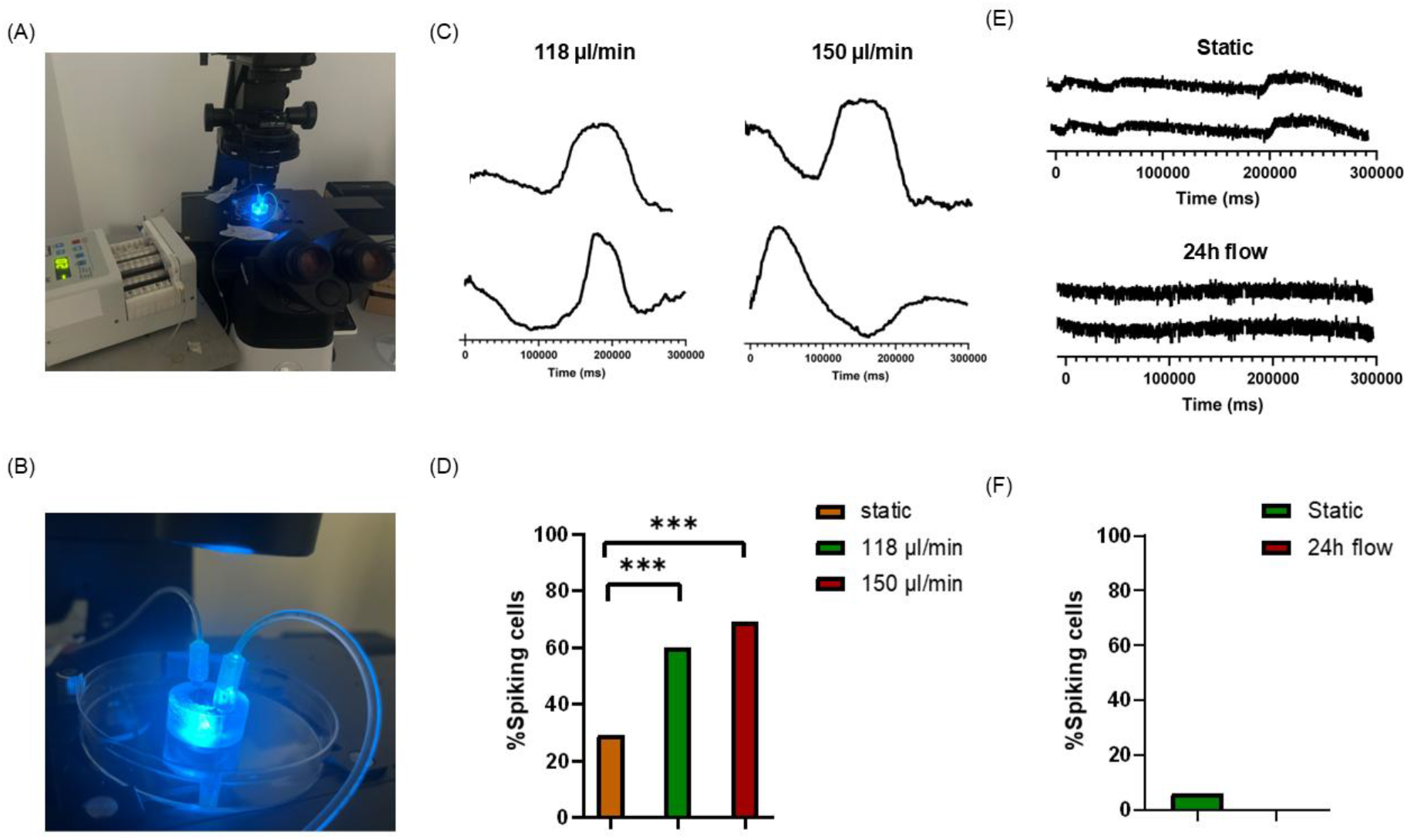
Live Ca^2+^ Imaging Measurements. (A) Epifluorescence microscope set up to perform live imaging with peristaltic pump. (B) Photograph of the microfluidic chip connected to the tubing on the epifluorescence microscope. (C) Calcium spikes of endothelial cells exposed to 118 µL/min (4.96x10^-2^ dyne/cm^2^) and 150 µL/min (6.30x10^-2^ dyne/cm^2^). (D) Analysis of the percentage of active endothelial cells in static, at 118 µL/min and 150 µL/min condition. (E) Calcium spikes of endothelial cells in static condition, and at 118 µL/min after 24h exposure. (F)Analysis of the percentage of active cells in static condition and after being exposed to 118 µL/min for 24h.

Cells were simultaneously visualized within the sample area in static condition (no flow), at 118 µL/min (4.96x10^-2^ dyne/cm^2^), and 150 µL/min (6.30x10^-2^ dyne/cm^2^), which are within the range of physiological conditions in humans. An average of 30 ± 10 fluorescence cells were imaged in each visual field within the sample area. In **Figure 3C** sample tracings of fluorescent recordings from activated endothelial cells are depicted for comparison between 118 µL/min and 150 µL/min. In live imaging recordings, repetitive and Ca^2+^ events were detected in 29% (44 out of 152 endothelial cells, n=4 visual field, n=4 independent series of culture) in static condition, 60% (92 out of 152 endothelial cells, n=4 visual field, n=4 independent series of culture) at 118 µL/min, 69% (105 out of 152 endothelial cells, n=4 visual field, n=4 independent series of culture) at 150 µL/min. The number of active cells was significantly (***p<0.001) higher in the two different flow-activated conditions compared to static cultures. No statistical difference was found between 118 µL/min and 150 µL/min (**Figure 3D**). These results suggest that endothelial cells modulate their calcium activity in response to rapid alteration of flow.

Hydrodynamic forces (e.g., shear stress), generated by blood flow act as fundamental mechanical that drives to vascular adaptation and endothelial homeostasis. Consistent with published results, endothelial cells have mechanosensors channel that drive to mechanotransduction when stimulated by forces (Mendoza et al., 2010; Nunez et al., 2024). Our live calcium imaging data demonstrate that acute and changes in shear stress generates a rapid mobilization of intracellular Ca^2+^, generating spikes in endothelial cells. Calcium functions as a key signaling molecule at the plasma membrane, modulating ion channel and receptor activity, and contributing to endothelial signaling and flow sensing (Gerhold and Schwartz, 2016).

These findings support the concept that dynamic mechanical forces can trigger Ca^2+^ mobilization, highlighting the central role of flow-mediated Ca^2+^ signaling in endothelial responsiveness

### 3.4 Long-Term Flow Abolishes Ca^2+^ Signaling

We next evaluated the effect of prolonged shear stress on endothelial cells by applying a continuous flow of 118 µL/min for 24 hours. Since no significant differences were observed between the flow rates of 118 µL/min and 150 µL/min, we selected 118 µL/min to minimize the risk of cell detachment under prolonged flow and to avoid introducing additional stress-related mechanisms that could affect cellular behavior. After that, cells were stained with Oregon Green 488 BAPTA-1 AM in fluidic condition and imaged as previously described. While endothelial cells showed flow-induced Ca^2+^ changes initially, these responses were absent following 24 hours of continuous shear stress. In **Figure 3E** sample recording of Ca^2+^ events in endothelial cells in static and after exposure to 118 µL/min flow rate. In these recordings, Ca^2+^ events were detected in 6% (5 out of 150 endothelial cells, n=4 visualized field, n=2 independent series of culture) in static condition, and 0% (0 out of 140 endothelial cells, n=4 visualized field, n=2 independent series of culture) at 118 µL/min for 24 h (**Figure 3F**).

Long-term exposure to flow (∼12-24 h) has been associated with adaptive behavior of endothelial cells (DeStefano et al., 2017). A plausible explanation is that prolonged shear stress induces feedback inhibition of mechanotransduction pathways, possibly through desensitization of shear-sensitive ion channels and homeostatic renormalization of intracellular Ca^2+^ levels (Davies, 1995). Likely, this flow-adapted state likely represents the physiological condition of endothelial cells, which are continuously exposed to blood flow and develop a protective phenotype (Chatterjee and Fisher, 2014).

### 3.5 Prolonged shear stress promotes endothelial junction formation

To quantify the barrier integrity in static and at 118 µL/min for 24h we performed immunocytochemistry to measure VE-Cadherin expression. To do so, HUVEC cells were immunostained for one of the main adherens-junction (AJ) protein (in red, **Fig. 4A**) and DAPI (in blue, **Fig 4**) for the nuclei.

**Figure 4.**
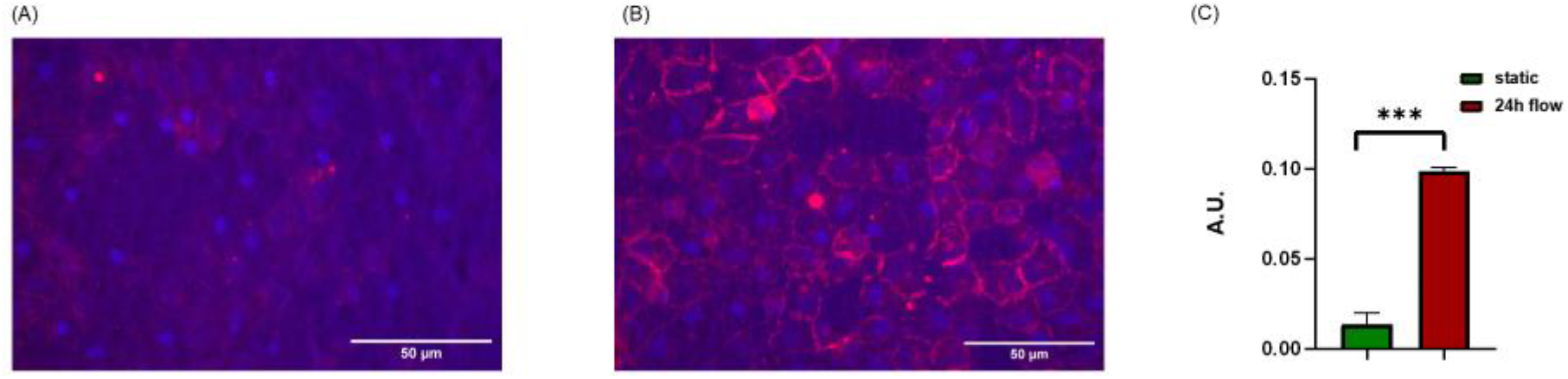
Adherens junctions’ increased expression in endothelial cells after 24h flow exposure. (A) Epifluorescence microscope image reconstruction of VE-Cadherin (red) and DAPI (blue) expression in endothelial cells in static condition. (B) Epifluorescence microscope image reconstruction of VE-Cadherin (red) and DAPI (blue) expression in endothelial cells after 24h at 118 µL/min (4.96x10^-2^ dyne/cm^2^). (C) Analysis of VE-Cadherin expression levels from the images presented in panel A-B, comparing static endothelial cells and 24-hour flow-exposed endothelial cells.

As sown in **Figure 4B**, the fluorescence expression of VE-Cadherin normalized on nuclei number, was significantly higher (0.098±0.002 A.U., n=6 visualized fields, n=2 independent series of cultures, ***p<0.001) after 24h flow-exposure compared to static condition (0.013 ± 0.006 A.U., n=6 visualized fields, n=2 independent series of cultures).

VE-Cadherin, a key component of endothelial junctions, plays a fundamental role in transmitting mechanical stress and activating intracellular signaling pathways that trigger biochemical cascades to modulate endothelial function (Chistiakov et al., 2017). When exposed to prolonged fluid shear stress (FSS; ∼12–24 h), endothelial cells exhibit junctional translocation toward the cell borders and alignment in the direction of flow, resulting in a tighter endothelial barrier (DeStefano et al., 2017; Franz et al., 2025; Noria et al., 1999).

It is well established that hemodynamic changes profoundly affect endothelial biology and morphology. However, the mechanisms linking altered flow to Ca^2+^ signaling remain poorly understood. Mechanosensitive channels, including the transient receptor potential (TRP) family and Piezo-1, have been identified in endothelial cells and modulate Ca^2+^ influx in response to mechanical stimuli such as shear stress (Cheng et al., 2023; Mendoza et al., 2010; Nunez et al., 2024). While calcium mobilization in response to shear stress is recognized as a key molecular signaling mechanism, little is known about live calcium responses in endothelial cells under long-term flow, primarily due to the lack of devices that allow both prolonged shear stress exposure and real-time imaging.

Previous studies have shown that endothelial cells undergo cytoskeletal rearrangements when exposed to flow, as confirmed by immunocytochemistry. Eskin et al. (1983) observed that endothelial cells align in the direction of flow, with this alignment becoming more pronounced after prolonged exposure. Consistent with these findings, Tovar-Lopez et al. (2019) characterized endothelial cell morphology by analyzing actin filament orientation, revealing alignment of cells in the direction of laminar shear stress.

Morphological alignment is not the only flow-dependent factor; shear stress also modulates junctional protein expression and other biological markers. VE-cadherin, a component of adherens junctions, is part of a mechanosensory complex with PECAM-1 that transmits mechanical signals to the cell, promoting adhesion and proliferation (Cheng et al., 2023; Simões-Faria et al., 2023). Similarly, Miao et al. (2005) reported that laminar flow is associated with sustained expression of adherens junction proteins such as VE-cadherin. More recently, Zhong et al. (2020) demonstrated that laminar flow stabilizes cell–cell interactions by reinforcing VE-cadherin-mediated junctions. Gene expression analyses have further supported this observation: Garcia-Polite et al. (2017) reported higher junctional protein expression in endothelial cells under laminar shear stress compared to static or disturbed flow conditions, while Rochfort et al. (2015) observed similar results after 24 hours of laminar shear stress. Our findings corroborate this mechanism, emphasizing the critical role of fluid flow in maintaining endothelial barrier integrity.

In this study, we developed a microfluidic platform that enabled investigation of how short- and long-term laminar shear stress modulates endothelial calcium signaling. Our platform incorporates a permeable membrane, which not only supports the structural organization of endothelial cells but also provides the opportunity to perform co-cultures with other cell types or to collect metabolites and soluble factors released under flow conditions. Our analysis revealed that endothelial cells exhibit robust flow-dependent Ca^2+^ responses during initial exposure; however, after 24 hours of continuous flow, these Ca^2+^ transients were no longer detectable, suggesting adaptation or desensitization of mechanosensitive pathways. Interestingly, prolonged shear stress was associated with enhanced expression of junctional proteins, indicating that chronic flow strengthens the endothelial barrier despite the loss of acute Ca^2+^ signaling. Previous studies have reported rapid intracellular calcium increases upon flow exposure, followed by adaptation under long-term laminar shear stress, resulting in a quiescent state that suppresses inflammatory activation (García-Polite et al., 2017; Kudo et al., 2003).

## Conclusions

In this study, we developed a low-cost, 3D-printed microfluidic platform that enables live imaging of endothelial cells under physiologically relevant shear stress. Our results demonstrate that rapid changes in flow activate endothelial mechanosensors, triggering Ca^2+^-mediated signaling, while prolonged flow induces structural remodeling of adherens junctions, enhancing barrier integrity. These findings highlight a temporal shift in endothelial mechanotransduction—from early calcium signaling to later junctional adaptation—and provide a versatile tool for studying vascular responses and cell–cell interactions under controlled flow conditions.

## Credit authorship contribution statement

LM designed and conceived the open-top platform and performed cell biology and immunofluorescence experiments and analysis. MH performed computational simulations. MT and LC contributed to chip design and development. RR conceived the study and the experimental design. LM, MH, MT, LC and RR wrote the manuscript.

## Acknowledgements

This work was supported by the European Union - NextGenerationEU within the framework of PNRR Mission 4 - Component 2 - Investment 1.1 under the Italian Ministry of University and Research (MUR) programme “PRIN 2022 PNRR” - grant number P2022TKL5T_001 Calinero - CUP: H53D23009070001.

## Data availability

The datasets generated for this study are available on request to the corresponding author.

